# Direct detection of RNA modifications and structure using single molecule nanopore sequencing

**DOI:** 10.1101/2020.05.31.126763

**Authors:** William Stephenson, Roham Razaghi, Steven Busan, Kevin M. Weeks, Winston Timp, Peter Smibert

## Abstract

Many methods exist to detect RNA modifications by short-read sequencing, relying on either antibody enrichment of transcripts bearing modified bases or mutational profiling approaches which require conversion to cDNA. Endogenous modifications are present on several major classes of RNA including tRNA, rRNA and mRNA and can modulate diverse biological processes such as genetic recoding, mRNA export and RNA folding. In addition, exogenous modifications can be introduced to RNA molecules to reveal RNA structure and dynamics. Limitations on read length and library size inherent in short-read-based methods dissociate modifications from their native context, preventing single molecule analysis and modification phasing. Here we demonstrate direct RNA nanopore sequencing to detect endogenous and exogenous RNA modifications over long sequence distance at the single molecule level. We demonstrate comprehensive detection of endogenous modifications in *E. coli* and *S. cerevisiae* ribosomal RNA (rRNA) using current signal deviations. Notably 2’-O-methyl (Nm) modifications generated a discernible shift in current signal and event level dwell times. We show that dwell times are mediated by the RNA motor protein which sits atop the nanopore. Further, we characterize a recently described small adduct-generating 2’-O-acylation reagent, acetylimidazole (AcIm) for exogenously labeling flexible nucleotides in RNA. Finally, we demonstrate the utility of AcIm for single molecule RNA structural probing using nanopore sequencing.

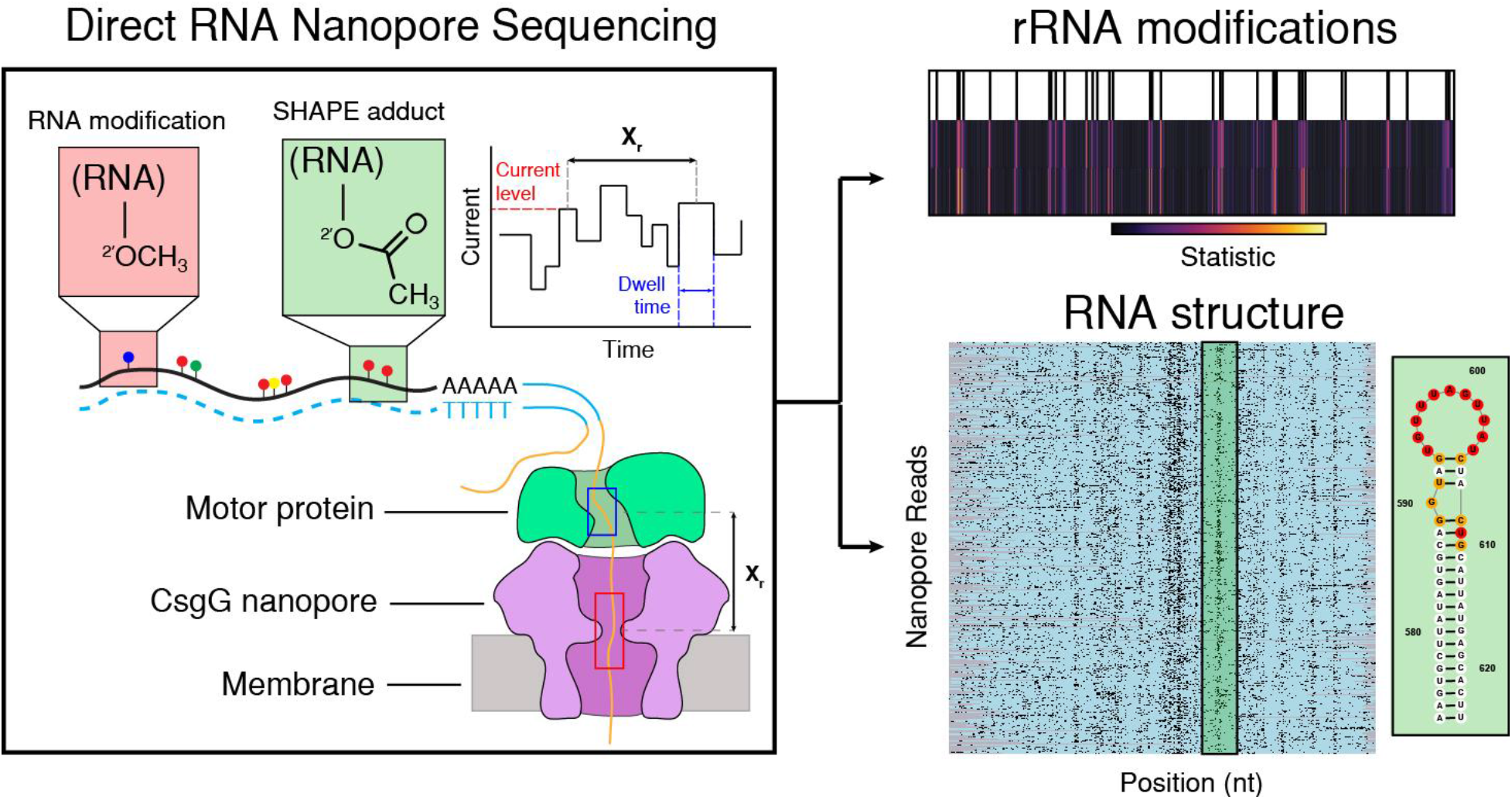
Graphical abstract.

## INTRODUCTION

To date over 100 distinct modifications of RNA have been identified, occurring on either the nucleobase or the ribose sugar. They have been shown to exhibit diverse effects on RNA including modulation of stability, translation efficiency, structural dynamics, export and translational recoding.(1–3) N6-methyladenosine (m^6^A), the most common mRNA modification, can be installed, read or removed by modification “writers”, “readers” and “erasers” respectively, suggesting a dynamic model of post-transcriptional gene regulation.(4, 5) Another modification, 2’-O-methylation (Nm) (methylation of the 2’ position of the ribose sugar), is found most prominently in the 5’ cap of mRNA (m^7^GpppNmNm) in eukaryotes and in ribosomal RNAs (rRNA). Recently, Nm modifications have also been detected within coding regions of mRNA (6) and have been demonstrated to have roles in tuning cognate tRNA selection during translation, thereby adjusting protein synthesis dynamics.(7) Pseudouridine (*Ψ*), often referred to as the fifth base due to its widespread inclusion in diverse classes of RNA, is generated by isomerization of uracil, thereby expanding hydrogen bonding and base pairing capabilities while also increasing base stacking propensity.(8, 9) The absence or reduction of RNA modifications has been shown to play a role in a variety of diseases including cancer (m^6^A), heart disease (m^6^A) (10), Treacher Collins syndrome (Nm) (11) and dyskeratosis congenita (*Ψ*).(12) Comprehensive detection and localization of RNA modifications within their native context would improve our understanding of RNA modification function and regulation, in addition to elucidating their role in disease.

A variety of methods exist for detection of specific RNA modifications, including pulldown of RNA using antibodies (immunoprecipitation) that specifically recognize modifications (e.g. m^6^A, 5mC) (e.g. MeRIP-seq, miCLIP).(13, 14) While immunoprecipitation based approaches can be applied transcriptome-wide, they only interrogate one modification at a time, do not give nucleotide resolution and are limited by the availability and effectiveness of antibodies. Furthermore, in conjunction with short-read sequencing, immunoprecipitation approaches require eventual fragmentation of RNA which destroys long-range information on phasing of modifications and complicates RNA isoform analysis. To determine the location of modifications with nucleotide resolution, reverse transcription (RT) and truncation approaches have also been employed. (6, 15–17) These approaches rely on manipulating either buffer conditions or the modified RNA directly, to encourage the RT enzyme to terminate prematurely upon encountering a base with either an endogenous modification or an adduct appended to a modification by a specific chemical reaction. While RT truncation-based methods provide localization information and estimates of modification quantification, they are complicated by RT-bias and cannot interrogate multiple modifications from the same single molecule. Mass spectrometry (MALDI or LC-MSMS) is currently the gold standard method for RNA modification identification and quantification as unique mass to charge ratios (m/z) and peaks provide information relating to precise chemical identities and abundances, however the method requires purified small RNA fragments for processing, which again precludes determination of information beyond the immediate primary sequence context of the site of modification.(18, 19)

Beyond endogenous RNA modifications, a variety of methods introduce exogenous modifications to assay certain aspects of RNA dynamics or structure. For example, metabolic labeling of nucleic acid involves the introduction of chemical analog nucleotides with moieties facilitating selective purification. In the case of RNA, ribonucleotide analogs such as 5-ethynyl uridine (5-EU) or 4-thio-uridine (s^4^U) can be incorporated into nascent transcripts, providing transcriptome-wide measurements of RNA stability, turnover and splicing dynamics.(20, 21) Additionally, a variety of chemical probes exist to assay RNA structure. These chemical probes target either the nucleobase to directly evaluate base pairing propensity (e.g. DMS, CMCT) or the ribose sugar to profile nucleotide flexibility (e.g. NAI, NMIA). The latter class of chemical probes are used in SHAPE (Selective 2’-Hydroxyl Acylation analyzed by Primer Extension) technologies and are capable of modifying all 4 ribonucleotides, in contrast to chemical probes such as CMCT and DMS which are most reactive to a subset of nucleobases under most conditions.(22–24) Location of modification with SHAPE reagents can be ascertained by performing RT and either measuring termination where the RT enzyme stalls at the modified nucleotide, or exploiting the propensity of RT in high Mn2+ to introduce mutations opposite sites of modification.

This latter approach, termed SHAPE-MaP (Selective 2’-Hydroxyl Acylation analyzed by Primer Extension and Mutational Profiling), (25) has been successful in determining single nucleotide resolution reactivity patterns transcriptome wide.(26) However, SHAPE technology relies on RT which produces shorter cDNA fragments when increasing the number of per-molecule SHAPE adducts. This limitation coupled with short-read sequencing precludes analysis of single molecule long-range structural information.(27)

Direct RNA nanopore sequencing has emerged as a promising long-read technology to detect sites of modification at the single molecule level. In the platform developed by Oxford Nanopore Technologies, currently the only commercially available direct RNA sequencing solution, RNA is translocated via a motor protein through a biological nanopore suspended in a membrane. (**Figure 1a**) As the RNA transits through the pore under voltage bias, the observed changes in picoampere ionic current are characteristic of the chemical identity (i.e. sequence) of a short k-mer (5 nucleotides) positioned at the pore constriction.(28, 29)

**Figure 1.**
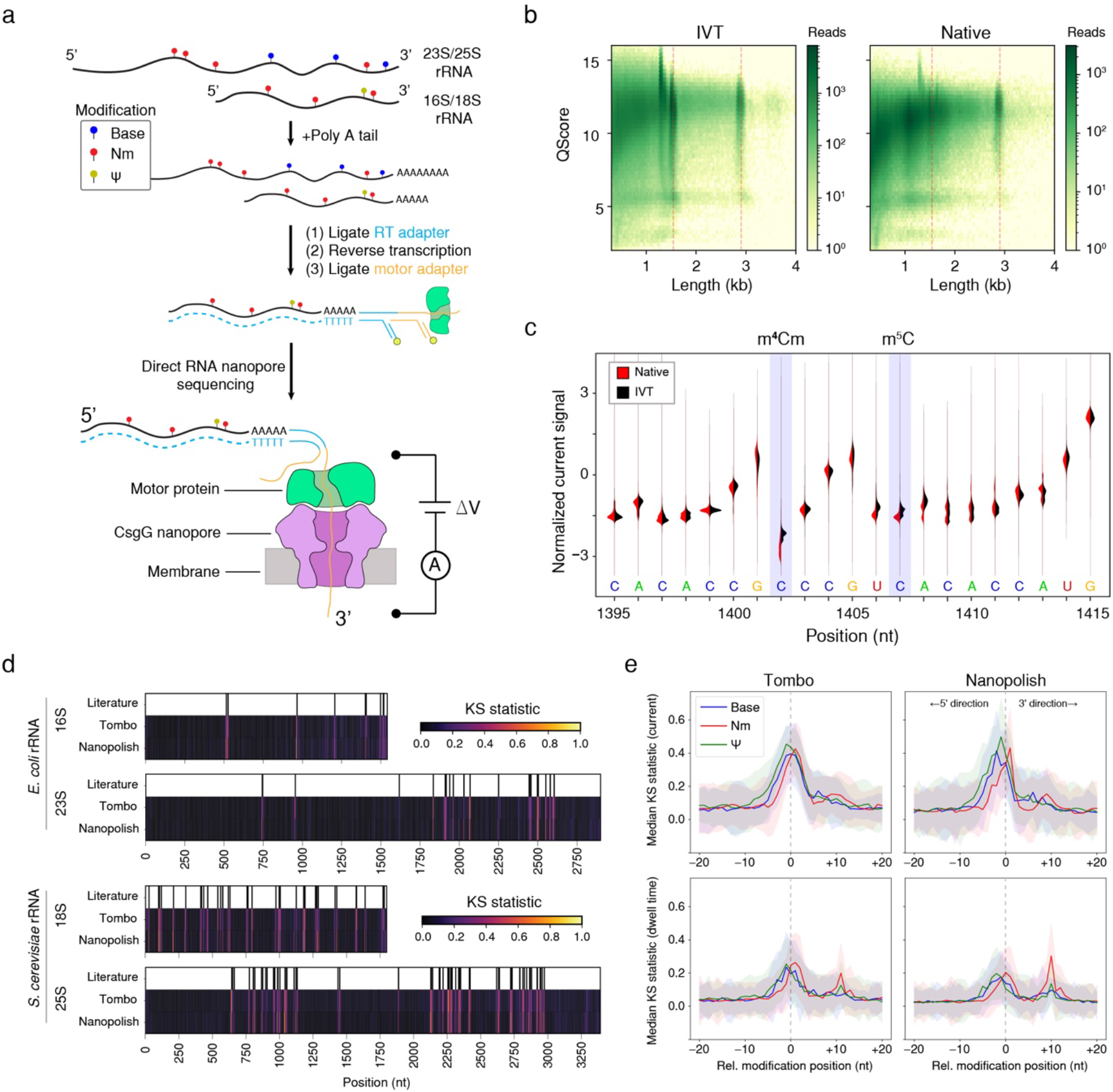
Direct RNA nanopore sequencing and modification detection. **a)** Direct RNA nanopore sequencing method. Ribosomal RNAs containing native modifications are poly(A) tailed before ligation of adapters and RT. Ionic current blockage events are characteristic of the k-mer sequence of RNA transiting through the pore constriction. **b)** Read quality heatmap for IVT rRNA (left) and native rRNA (right) from *E. coli*. Dashed red lines indicated the expected lengths for 16S rRNA (1.5 kb) and 23S rRNA (2.9 kb) **c)** Normalized native (red) and IVT (black) current signal alignment for 16S rRNA from *E. coli* spanning positions 1395 - 1415 performed using Tombo. Sites of known modifications within this window are highlighted in blue. **d)** Positional Kolmogorov-Smirnov (KS) statistical testing across rRNA from *E. coli* and *S. cerevisiae* using both Tombo and Nanopolish. Known modification positions (Literature) (37, 38) are indicated above as black lines. **e)** Median current and dwell KS statistic profiles separated by modification type (Base - blue, 2’ -O-methyl - red and pseudouridine - green) aligned by modification position from both Tombo and Nanopolish. Colored shaded regions represent the standard deviation of the KS statistic.

Here we use the raw current signal from nanopore sequencing to detect the presence of RNA modifications, agnostic to the chemical nature of the modification. For well-characterized endogenous modifications in rRNA, we reliably detect current perturbations to within ± 2 nt of the expected positions. In addition, we describe modification-dependent signals in the time domain, providing a complementary dimension of information that may be incorporated with current perturbations for *de novo* identification of nucleotide modification classes. We also investigated our ability to detect exogenous RNA modifications to enable structural profiling. First, we tested common SHAPE reagents for compatibility with nanopore sequencing. We observe that commonly used SHAPE reagents (e.g. NAI, NMIA) are not suitable for nanopore sequencing likely due to unfavorable interactions between the bulky 2’-O-aryl adduct and the motor protein. This motivated a search for an alternative SHAPE reagent with a smaller adduct which would be more compatible with the nanopore sequencing platform. By screening a variety of acylimidazole candidate probes we identified acetylimidazole (AcIm) as having favorable reactivity with RNA. We demonstrate that AcIm is a cell-permeable SHAPE-MaP reagent by verifying the structure of extracted ribosomes and nuclear located U1 snRNA. Finally, we demonstrate that AcIm is suitable for single molecule RNA structural profiling on the nanopore platform, a technique we call nanoSHAPE (Selective 2’-Hydroxyl Acylation analyzed by nanopore sequencing). Investigating the structure of the pri-miR-17∼92 microRNA cluster we detect more modification sites per RNA molecule and over longer distances compared to SHAPE-MaP experiments. We find that structural predictions of pri-miR-17∼92 constrained by nanoSHAPE and SHAPE-MaP agree well, suggesting nanoSHAPE is a promising method to detect RNA structure at the single molecule level.

## MATERIALS AND METHODS

### rRNA extraction (E. coli and S. cerevisiae)

*E. coli* (K-12 MG1655 and EDCM648-d ΔmraW54/ΔrsmH) cells were grown in an overnight culture at 37 °C in 10 mL of freshly prepared Luria Bertani broth (LB) or 10 mL M9 minimal salts. 75 mL of pre-warmed media was inoculated with 1 mL of overnight culture. *E. coli* cells were grown to OD_600_ = 0.5, typically over 3 - 4 hours at 37 °C. 25 mL of *E. coli* cells were pelleted at 3280 .g at 4 °C for 12 minutes. Cells were lysed in 16.5 mL lysis buffer (15mM Tris pH 8, 450 mM sucrose, 8 mM EDTA, 0.4 mg/ml lysozyme) for 5 minutes at room temperature then 10 minutes at 4 °C. The pellet was collected at 3280 °g for 5 minutes and then resuspended in 2 mL proteinase K buffer (50 mM HEPES, 200 mM NaCl, 5 mM MgCl_2_, 1.5% SDS, 0.2 mg/mL Proteinase K). The solution was vortexed for 10 seconds and incubated at room temperature for 5 minutes, and then 4 °C for 10 minutes. Nucleic acids were extracted twice with 1 volume of phenol:chloroform:isoamyl alcohol (25:24:1) followed by two subsequent chloroform extractions prior to ethanol precipitation and resuspension in 88 μl RNAse-free water. Purified nucleic acids were treated with Turbo DNAse (10 μl Turbo DNAse buffer [10.] and 2 μl Turbo DNAse) at 37 °C for 1 hour. Finally, RNA was purified with 0.8X vol AmpureXP beads. For *S. cerevisiae* (S288C), colonies were picked from an agar plate and incubated at 30 °C in ∼7 mL of YPD broth for two days. RNA purification was carried out using the YeaStar RNA kit (Zymo Research) according to instructions.

### Generation of rRNA IVT controls

gDNA was extracted from *E. coli* using the method described for RNA above up to and including the ethanol precipitation step. After resuspension in 88 μl RNAse-free water, purified nucleic acids were treated with 1-5 μl 1 mg/ml RNaseA (Qiagen) for 45 minutes at 37 °C to degrade RNA. gDNA was then purified with 0.5X SPRI or subsequent ethanol precipitation. gDNA was purified from *S. cerevisiae* using the YeaStar Genomic DNA kit (Zymo Research) according to instructions. Primers for amplifying rDNA amplicons, which include a T7 transcription promoter for subsequent in vitro transcription (IVT) are detailed in Supplementary Table 1. Amplicons were generated by PCR using Kapa HiFi DNA polymerase and purified by SPRI. T7 transcription templates were transcribed using HiScribe T7 Quick High Yield RNA Synthesis Kit (New England Biolabs). Reactions were cleaned up using the MEGAClear Transcription Clean-up Kit (Invitrogen) before nanopore sequencing library preparation.

### Poly(A) tailing of RNA

Oxford Nanopore Technologies direct RNA sequencing requires a poly(A) tail for first adapter ligation. Both rRNA and pri-miR-17∼92 samples were poly(A) tailed using *E. coli* Poly(A) Polymerase (New England Biolabs) according to manufacturer’s instructions.

### Nanopore library preparation

Direct RNA sequencing was performed using the Oxford Nanopore Technologies kit (SQK-RNA002) with the RCS control RNA. Sequencing was performed on the MinION device using either standard flowcells (FLO-MIN106D) for rRNA experiments or flongle flowcells (FLO-FLG001) for pri-miR-17∼92 experiments. Sequencing was carried out until the number of active nanopores dropped below 5% of the initial total number of pores, typically 12-36 hours.

### Nanopore data processing

Multi-fast5 reads were basecalled using guppy (v3.1.5). Base called multi-fast5 reads were then converted to single read fast5s using the Oxford Nanopore Technologies API, ont_fast5 (v1.0.1). Fastqs were mapped to their respective transcriptomes for *E. coli* (NC_000913.3.fa) and *S. cerevisiae* (R1-1-1_19960731.fsa) using minimap2 (v2.11).

### Nanopolish and Tombo analysis of data

Tombo (v1.5.1) and Nanopolish (v0.11.1) were both used to detect native modifications in rRNA datasets as well as detect modifications deposited from SHAPE reagents. Comparisons were performed between native and IVT samples for the rRNA datasets and between modified (at indicated concentrations) and unmodified samples for pri-miR-17∼92. Nanopolish eventalign module was used to align current intensities and dwell times to reference sequences. Kolmogorov–Smirnov (KS) statistical testing was performed in order to detect modified nucleotides. Using Tombo, raw signal squiggles were assigned to reference sequences using *resquiggle*. Next, modified base detection was carried out using the *detect_modifications model_sample_compare* method. Per-read statistical testing (AcIm modified RNA) was performed with a ± 1 nucleotide Fisher’s method context adjustment. The requisite text output was obtained using *text_output browser_files* method. Reactivity profiles from Tombo per-read statistical testing were further adjusted using Benjamini-Hochberg procedure for multiple testing. Adjusted per-read reactivity profiles were used to calculate percentage modification per genomic position. This percentage profile was then normalized using the normalization procedure described in SHAPE-MaP method (30). Single molecule positional current, standard deviation of current, and dwell time data were extracted as numpy arrays directly from single read fast5 data using custom written python scripts.

### Reagents

All standard laboratory reagents, including AcIm, were purchased from Sigma-Aldrich, with the exception of NMIA purchased from Thermo Fisher Scientific. NAI was synthesized from 2-methylnicotinic acid and 1,1′-carbonyldiimidazole, as described. (31)

### AcIm hydrolysis

AcIm hydrolysis was tracked at 37 °C by time resolved UV absorbance using a Nanodrop 2000 spectrophotometer in [1.] modification buffer (100 mM HEPES pH 8.0, 100 mM NaCl, 10mM MgCl_2_) every 2 minutes for 40 minutes. Imidazole spectra were collected every 2 minutes in [1.] modification buffer for 40 minutes.

### SHAPE-MaP on E. coli 16S and 23S rRNAs

Extracted RNA was treated with NAI [100 mM final], AcIm [100 mM final], NMIA [13 mM final] or DMSO (unmodified control). All SHAPE-MaP experiments were performed with 10% volume fraction of DMSO. Modification was carried out at 37 °C for 3 half-lives of the chemical probe used. For mutational profiling RT, 1 μl of nonamer primer [200 ng/μl or 2μM] was added to 1-3 μg of rRNA in 10 μl nuclease free water. The samples were incubated at 65 °C for 5 minutes then cooled on ice. 8 μl of [2.5×] MaP buffer (125 mM Tris pH 8.0, 187.5 mM KCl, 15 mM MnCl_2_, 25 mM DTT, and 1.25 mM dNTPs) was added and incubated at 42 °C for 2 minutes. 1 μl of SuperScript II reverse transcriptase was added and mixed well before incubating the reaction at 42 °C for 2-3 hours, and then at 70 °C to inactivate the polymerase. cDNA was exchanged into water using G-50 columns (GE Life Sciences) the volume increased to 68 μl using nuclease free water. Second strand synthesis was carried out (Second Strand Synthesis Enzyme mix; New England Biolabs) and the dsDNA was used to generate a library for sequencing on an Illumina MiSeq, as described.(30)

### SHAPE-MaP on U1 snRNA

LCL (Coriell) cells were cultured in RPMI 1640 media supplemented with 2 mM L-glutamine and 15% fetal bovine serum. LCL cells were collected and resuspended in fresh media prior to addition to either 100 μl of 2 M AcIm [200 mM final] or 100 μl of 2 M NAI [200 mM final] in DMSO for 4-5 minutes. RNA was extracted and SHAPE-MaP was performed as described.(32) RT was performed using a U1 snRNA-specific primer (5’-CAGGG GAAAG CGCGA AC-3’)

### RNA modification (pri-miR-17∼92)

In order to protect the 3’-OH of pri-miR-17∼92 RNA from modification with acylating reagents, the terminal 3’ nucleotide was oxidized followed by a beta-elimination reaction to remove the terminal nucleotide leaving a terminal phosphate. Then RNA modification was carried out prior to dephosphorylation and nanopore library preparation. Briefly, pri-miR-17∼92 was incubated at 37 °C for 30 minutes with shaking in oxidation buffer (NaIO_4_ [20mM], Lysine-HCl [200mM] pH 8.5, final volume: 40 μl). The reaction was quenched with 2 μl of ethylene glycol then purified using 1. SPRI, eluting into beta-elimination buffer (Sodium borate [33.75mM], boric acid [50mM], pH 9.5) incubating at 45 °C for 45 minutes. RNA was again purified by 1. SPRI. 1-2.5 μg of IVT pri-miR-17∼92 RNA was diluted into 7 μl water and heated to 95 °C for 2 min and immediately placed on ice (2 min). 6μl of folding buffer [3.3.] (333 mM Tris-HCl pH 8.0, 333 mM NaCl and 33 mM MgCl2) and 5 μl HEPES pH 8.0 [200 mM] were added and the RNA was allowed to fold for 20 min at 37 °C. 2 μl of DMSO (control) or SHAPE reagent (NAI or AcIm) were added to a new tube, then folded RNA was added and mixed by pipetting. Modification was carried out for at least 3 half-lives at 37 °C. RNA was then dephosphorylated by adding 22 μl RNAse-free water, 5 μl Antarctic Phosphatase reaction buffer [10°], 2 μl Antarctic Phosphatase [5k U/mL] (NEB) and 1 μl RNase inhibitor and incubating at 37 °C for 30 minutes with shaking. The phosphatase was inactivated by incubating the reaction at 65 °C for 5 minutes. Finally the RNA was purified using 1. SPRI prior to poly(A) tailing.

### RNA structure modeling

Centroid free energy structures and energies were obtained using the RNAfold (v2.4.13) (Vienna) web server. (http://rna.tbi.univie.ac.at//cgi-bin/RNAWebSuite/RNAfold.cgi) Options were to avoid isolated base pairs and temperature = 37 °C. R-chie (33) was used for displaying base pairing (arc) of centroid structures. The RNAstructure (v6.2) software suite (34) was used for partition function calculation and associated dot plot visualization. The following options were used for partition function calculation: maximum percent energy difference = 10%, maximum number of structures = 50, window size = 3, temperature = 37 °C.

## RESULTS

### Identification of specific modifications at defined locations within 16S rRNA

To assess the capability of the nanopore platform to detect RNA modifications directly on rRNA we generated in vitro transcribed (IVT) controls completely devoid of modifications for the small and large ribosomal subunits through direct amplification of rDNA from *E. coli* and *S. cerevisiae* (**Supplementary Figure 1a,b**). IVT controls, which were sequenced independently, allow direct comparison to the natively modified rRNA facilitating modification detection. Both the IVT controls and native experiments exhibited reads at the expected rRNA lengths for *E. coli* (16S: 1.5 kb, 23S: 2.9 kb) (**Figure 1b**) and *S. cerevisiae* (18S: 1.8 kb, 25S: 3.4 kb) (**Supplementary Figure 1c**). Notably, median QScores for full length molecules were lower for native samples compared to IVT controls (16S: −0.53, 23S: −0.31, t-test p < 0.05) (**Supplementary Figure 1d**), likely reflecting the effect that high levels and densities of modifications have on read quality using the current iteration of modification un-aware base calling software. (**Materials and methods**) Generally, coverage was lower towards the 5’-end for all samples including IVT controls, (**Supplementary Figure 2**) as expected from the configuration of direct RNA nanopore sequencing which translocates RNA in the 3’ to 5’ direction.

To investigate the presence of modifications, reads were processed using both Tombo (35) and Nanopolish (36), which perform raw signal level to sequence alignment. (**Materials and methods**) As an example, we highlight a region of current signal mapped with Tombo on 16S rRNA in *E. coli* containing two modifications in close proximity: N4,2’-O-dimethylcytosine (m^4^Cm) at position 1402 and 5-methylcytosine (m^5^C) at position 1407. (**Figure 1c**) We observed a clear deviation in the current signal distribution of the native sample at position 1402 which contains a base methylation on the 4 position of cytosine in addition to methylation of the 2’-hydroxyl (2’-O-methyl or Nm). Current signal deviation due to the chemically distinct m^5^C modification at position 1407 was less pronounced and spread over 3 positions, 1406 - 1408, (**Figure 1c**) highlighting the often distributed nature of complex nanopore signals in the k-mer context.

### Comprehensive rRNA modification detection

We next set out to examine detection of all known modifications (**Supplementary data 1**) in the small and large subunit rRNA of both *E. coli* and *S. cerevisiae*.(37, 38) We performed non-parametric Kolmogorov-Smirnov testing (KS) across all positions for current and dwell time from raw signal aligned data using both Tombo and Nanopolish. (**Materials and Methods**) (**Figure 1d**) Peaks in the KS statistic profile indicate distributional differences between the IVT unmodified control sample and the corresponding natively modified samples. The KS statistic for the current signal was strongly correlated between Tombo and Nanopolish (Spearman’s rank order correlation, rho = 0.49 - 0.64) however the dwell time KS statistic profiles were only moderately correlated (Spearman’s rho r = 0.31 - 0.41). Generally, KS statistic peaks for current are observed within +/−2 nt of the known modifications. To obtain a general assessment of modification class features we collated all RNA modifications across the small and large ribosomal subunits from both *E. coli* and *S. cerevisiae*, aligning them by their known modification position and considered the aggregate median KS statistic profile across modification class (Base, Nm, or *Ψ*). (**Figure 1e**) The median KS statistic for current of all modification types had an appreciable signal at the site of modification (pore constriction) as expected (**Figure 1e top row**). Regarding dwell time, the median KS statistic profile exhibited two peaks for Nm and pseudouridine modifications but not for base modifications. (**Figure 1e bottom row**) The primary peak occurred at the nanopore constriction (relative modification position 0) however, secondary peaks were observed approximately 10 nucleotides in the 3’ direction (i.e. prior to the modification reaching the pore constriction) with Nm modifications exhibiting a larger median KS statistic than *Ψ*. We surmise that the 10 nucleotide distance is the ‘registration distance’ (Xr) from the pore constriction (where k-mer currents are measured) to the motor protein which sits atop the nanopore. The Xr identified here is consistent with previous measurements of registration distance in DNA nanopore sequencing experiments using a more terminally located pore constriction in MspA (Xr ∼ 20nt) along with a different motor protein.(39) In the R9.4.1 version nanopore system used here, a CsgG pore is used which has a centrally located pore constriction suggesting a smaller Xr.(28)

Interestingly, these observations suggest that at least in some sequence contexts the motor protein kinetics are sensitive to Nm modifications and to a lesser degree, *Ψ* modifications. We note that 2’-O-methylation confers approximately −0.2 kcal/mol of stacking free energy to ssRNA and induces the C3’-endo conformation of RNA (40) possibly perturbing translocation times due to steric or chemical interactions with particular amino acid residues of the motor protein. Interestingly, all types of modifications exhibit a median KS statistic peak for dwell time above background at relative modification position 0, indicating that transit times through the nanopore constriction itself are also perturbed by many of the modifications studied here.

### RNA Translocation rate is sensitive to nucleotide modifications and sequence composition

We next investigated the dwell time as a function of position to ascertain whether modifications and or sequence content perturb dwell times. Comparison of dwell times for both native and IVT samples at a distance of +Xr from selected Nm sites within 16S (1402 m^4^Cm), 23S (2552 Um), 18S (1428 Gm) and 25S (2220 Am) rRNA revealed a significant increase (Mann-Whitney U test). (**Figure 2a**) To assess how much of motor protein pausing is due to sequence context removed from the presence of modifications, we explored sequence similarities of the top 1% of ranked dwell times from all IVT samples in the region spanning the entire sequencing complex including upstream of the motor protein to the exit of the nanopore. We observed a very strong guanosine enrichment approximately 9 - 11 nt (motor protein active site ‘tri-mer’) upstream of the pore constriction (**Figure 2b**), consistent with the registration distance Xr = 10 nt, and with increased dwell time observations for Nm modifications. More broadly, we quantified the fraction of each nucleotide in the motor protein active site tri-mer across dwell time percentiles for all IVT samples, and observed overrepresentation of guanosine in the highest percentiles. (**Figure 2c**) Collectively these data indicate that even absent modifications, the motor protein used in direct RNA nanopore sequencing experiments has a tendency to pause on guanosine rich sequences (e.g. polyG tracts). This has been previously observed in the context of single-molecule picometer resolution nanopore tweezers (SPRNT) experiments on DNA which employed a Hel308 based translocation mechanism.(41) Further, this may be a general feature of enzyme-based translocation along polyG tracts due to steric hindrance and or higher than average single stranded stacking energies.(42)

**Figure 2.**
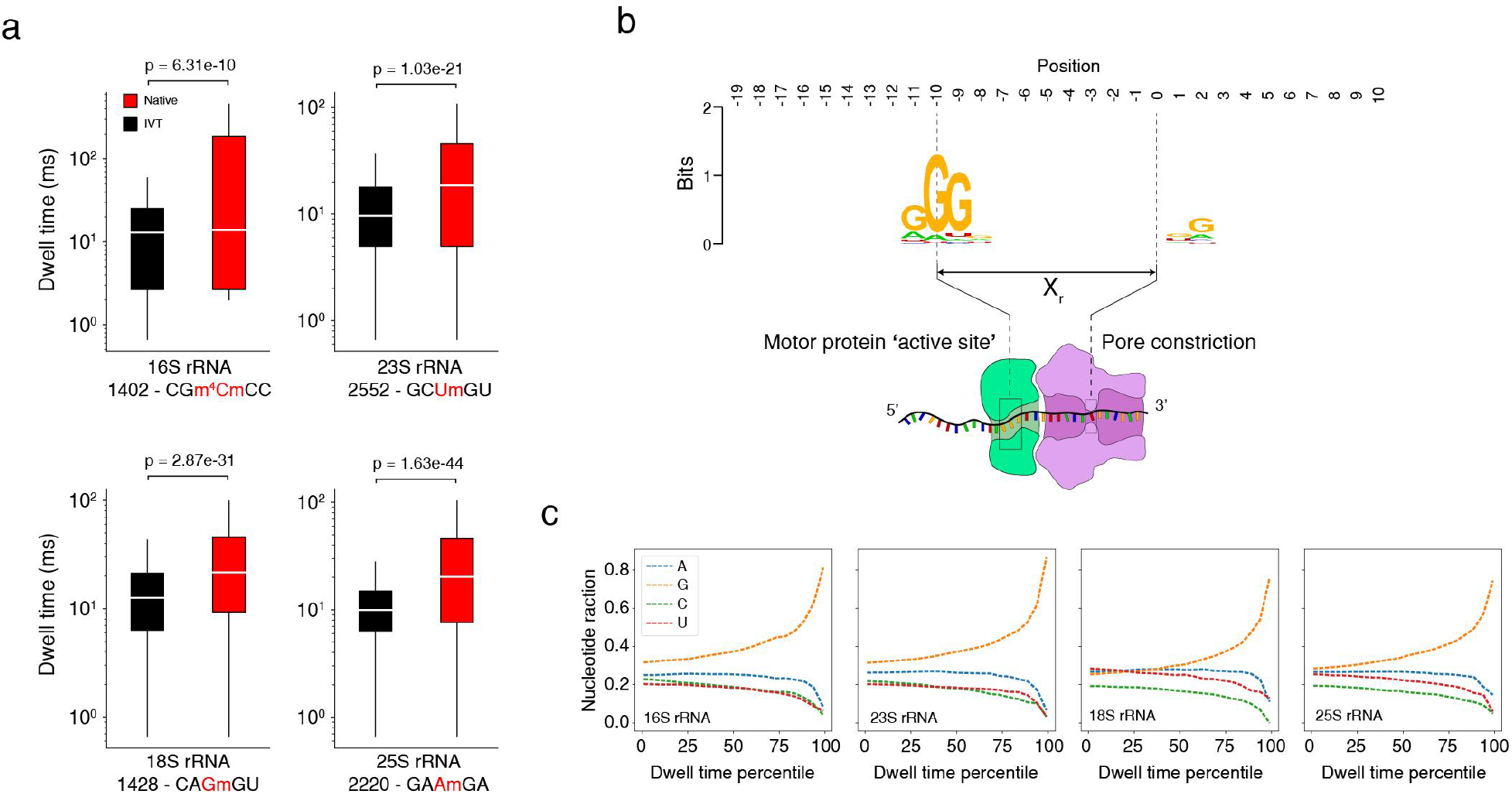
Dwell time dependence on modifications and sequence. **a)** Dwell time comparison of native (red) and IVT (black) samples (Mann-Whitney U test, n = 1000 reads) at selected Nm modification sites in 16S, 23S and 18S, 25S from *E. coli* and *S. cerevisiae* respectively. The k-mer is indicated below the x-axis along with the position of the modification (red). Dwell times are from +Xr from the centered modification site k-mer. **b)** Sequence motif from the top 1% of dwell times (IVT samples only, 16S, 23S, 18S, and 25S) spanning a 30 nucleotide window encompassing the entire biomolecular sequencing complex **c)** Trimer (positions −9,−10, and −11) nucleotide fraction from IVT samples as a function of dwell time percentile. The highest dwell time percentiles are enriched for guanosine within the trimer.

### 1-acetylimidazole is a small adduct-generating, cell-permeable, SHAPE-MaP reagent

After demonstrating detection of endogenous Nm modifications in rRNA, we hypothesized that we could detect 2’-O-adducts resulting from exposure of folded RNAs to electrophilic SHAPE reagents, enabling us to interrogate RNA structure using nanopore sequencing. We initially attempted nanopore sequencing using RNA that had been modified with established SHAPE reagents (NAI, 1M7 and NMIA). These experiments resulted in low numbers of sequenced molecules, few full length reads, and poor alignment accuracy as compared to an unmodified control sample (**Supplementary Figure 3**). We speculate that the observed inefficient and incomplete translocation was due to the presence of large 2’-O-aryl adducts, that result from reaction of RNA with these SHAPE reagents. We therefore searched for a nanopore compatible reagent which would produce a smaller, chemically similar adduct to Nm modifications which would be amenable to nanopore sequencing. Ideally the reagent would retain high and relatively uniform reactivity levels at the 2′-hydroxyl position across the four canonical ribonucleotides. We explored the suitability of AcIm (43), along with 4 other carbonyl-imidazole candidates, for structure-selective 2′-O-acylation. We detected covalent adduct formation for the expected positive control (NAI) and for AcIm (**Supplementary Figure 4**).

The proposed reaction of AcIm with the 2′-hydroxyl of RNA (**Figure 3a**) results in an acetyl adduct that is essentially the most compact adduct possible via a carbonyl electrophilic reagent. The relative rate of a SHAPE reagent reaction with the 2’-hydroxyl of RNA is mirrored by its rate of reaction with water.(22) Therefore we investigated the timescale of AcIm reactivity by monitoring the absorbance of AcIm in reaction buffer; imidazole was monitored as a control. For AcIm, an absorbance signal, centered at 250 nm, decayed via a single exponential, corresponding to a half-life of approximately 3 minutes at 37 °C (**Figure 3b**), consistent with prior measurements of N-acetyl-imidazole hydrolysis.(43, 44) The AcIm spectrum decays to that consistent with imidazole, supporting hydrolysis of AcIm into imidazole and unreactive, non-absorbing acetate. To assess AcIm reactivity with RNA, we extracted RNA from *E. coli* and treated total RNA with either NAI [100 mM], NMIA [13 mM], AcIm [100 mM], or DMSO (unmodified control). We then performed mutational profiling (SHAPE-MaP) and aligned the resulting cDNAs to the 16S and 23S rRNAs and compared reactivities across reagents (**Figure 3c**). Reactivity profiles from all three reagents showed high correlation across both rRNAs, indicating that AcIm is a *bona fide* SHAPE reagent. AcIm reacted broadly with all four canonical RNA nucleotides and showed good discrimination for reaction with single-stranded and conformationally flexible nucleotides (**Supplementary Figures 5 and 6**). To assess cell permeability, we performed MaP on U1 snRNA from a lymphoblastoid cell line exposed to either AcIm or NAI. Reactivity profiles from the reagents were strongly correlated (Spearman’s rank order correlation, rho = 0.72) (**Figure 3d**), corroborate the known secondary structure of U1 snRNA (**Figure 3e**), and are consistent with in-cell reactivity patterns of widely used SHAPE reagents. In sum, AcIm is a cell-permeable SHAPE reagent that reacts with all four ribonucleotides and generates SHAPE-MaP reactivity profiles consistent with known reagents and is useful for RNA structure modeling.

**Figure 3.**
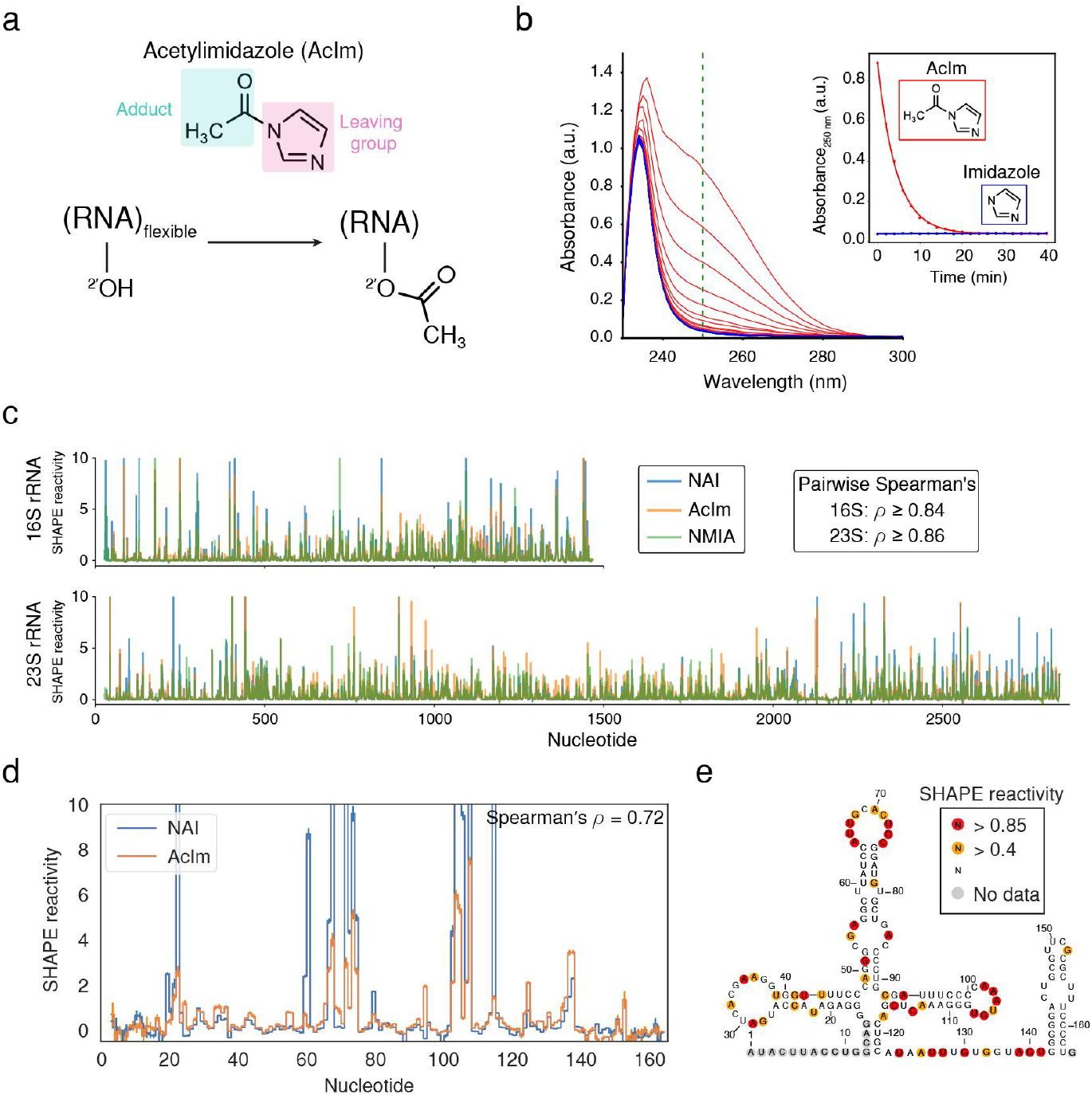
Acetylimidazole is a cell-permeable SHAPE-MaP reagent. **a)** Acylation of RNA with acetylimidazole (AcIm). **b)** Time resolved hydrolysis of acetylimidazole analyzed by changes in UV absorbance at 37 °C. **c)** SHAPE-MaP reactivity profiles using NAI, AcIm, or NMIA for *E. coli* 16S and 23S rRNA. **d)** In-cell SHAPE-MaP probing of U1 snRNA with AcIm and NAI. **e)** Secondary structure of U1 snRNA overlaid with reactivity from AcIm-based in-cell SHAPE-MaP.

### nanoSHAPE: Direct RNA nanopore sequencing of AcIm modified RNA

We next assessed whether AcIm chemical probing could be used to guide RNA secondary structure modeling based on single molecule direct RNA nanopore sequencing, a method we call nanoSHAPE. We focused on an in vitro transcribed pri-miRNA transcript of the miR-17∼92 cluster, which spans 951 nucleotides and folds to form a series of hairpin structures.(45, 46) Pri-miR-17∼92 is stable and predicted to form a moderately structured (Predicted ΔG_centroid_ = −298.80 kcal/mol) complex with a comparably low number of near-energy suboptimal conformations given its length (ensemble diversity, ED = 152.9) making it a suitable substrate for in vitro folding studies. We verified the structure of pri-miR-17∼92 by performing SHAPE-MaP on in vitro transcribed and folded pri-miR-17∼92 using both AcIm and NAI. (**Supplementary Figure 7a**) SHAPE-MaP reactivity was highly correlated between AcIm and NAI (Spearman’s rho = 0.78). Furthermore, the structure obtained from secondary structure prediction using RNAfold constrained with the AcIm SHAPE-MaP reactivity profile produced a centroid structure consistent with the known pri-miR hairpins comprising the 17∼92 cluster.(45) (**Supplementary Figure 7b**) These experiments indicate that both chemical probes are suitable for investigating this structure in an in vitro context and additionally provide a control structure for comparison with nanoSHAPE. Next, to assess the compatibility of AcIm with nanopore sequencing we performed a series of direct RNA nanopore sequencing experiments using 8 individual flongle flow cells on pri-miR-17∼92 (3’- oxidized) modified with 0mM (unmodified control), 5mM, 20mM, 50mM, 75mM, 100mM, 150mM and 200mM final concentrations of AcIm. (**Figure 4a**) Read quality and the number of full length reads noticeably decreased with increasing concentration of AcIm, decreasing the resultant percentage of successfully aligned reads using Tombo. (**Supplementary Figure 8a, d, e and Supplementary Table 1**) The coverage was higher at the 3’ end for all concentrations, again consistent with the known read direction of the RNA through the nanopore (3’-> 5’) (**Supplementary Figure 8b**) The coverage, fraction of full length reads and aligned read percentage was inadequate for the highest AcIm concentration (200mM) so this condition was excluded from further analysis.

**Figure 4.**
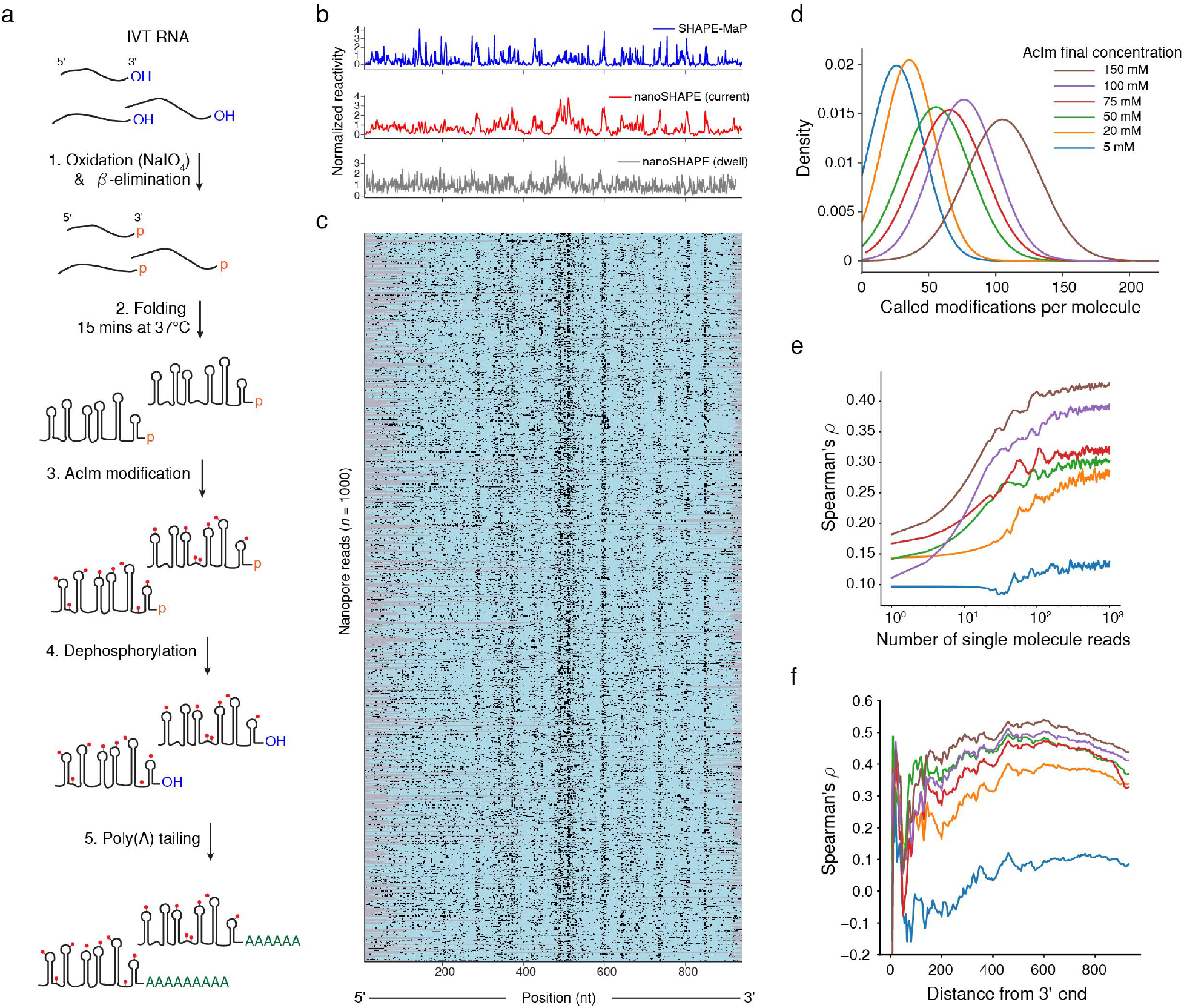
Direct RNA structural probing of pri-miR-17∼92 using nanopore sequencing. **a)** nanoSHAPE protocol: IVT RNAs with 3’-hydroxyl are oxidized prior to ***β***-elimination to remove the terminal nucleotide and create a 3’-phosphate RNA (one nucleotide shorter than the starting RNA). RNAs are folded and modified with AcIm prior to dephosphorylation and poly(A) tailing. Poly(A) tailed RNA is used as input for nanopore sequencing. The control (unmodified) sample was processed identically without the AcIm modification step. **b)** Normalized SHAPE-MaP reactivity (blue), normalized nanoSHAPE reactivity (current) (red), normalized nanoSHAPE reactivity (dwell time) (grey) for pri-miR-17∼92 (top). **c)** Heatmap of 1000 AcIm modified (150mM) pri-miR-17∼92 nanopore reads. Modifications are determined by per base Student’s t-test using Fisher’s method context of ± 1. Per-base p-values are corrected using the Benjamini-Hochberg procedure and binarized. Modifications are shown in black, unmodified positions are shown in teal, unmapped regions are shown in grey. **d)** Kernel density estimate (KDE) of the number of called modifications (2’-O-acetyl adducts) per pri-miR-17∼92 molecule across the AcIm concentrations tested. **e)** Spearman’s rank order correlation (rho) with SHAPE-MaP as a function of the number of contributing pri-miR-17∼92 molecules across the AcIm concentrations tested. **f)** Spearman’s rank order correlation (rho) with SHAPE-MaP as a function of the distance from the 3’-end of the pri-miR-17∼92 RNA.

Next, we performed KS statistical testing for current and dwell time distributions across all AcIm concentrations compared to the unmodified control. (**Supplementary Figure 9a**) KS peaks in current were observed in all profiles consistent with locations of single stranded regions of pri-miR-17∼92. We performed Spearman’s rank order correlation (rho) for both KS of current and KS of dwell time (shifted by Xr) against the AcIm SHAPE-MaP profile. (**Supplementary Figure 9b**) The rank correlation was greatest for current, and maximized at 150mM (Current rho = 0.51, dwell time rho = 0.28). In order to assess the capability of single molecule based reconstruction of reactivity profiles we performed per-read statistical testing and normalization for every nucleotide position within 1000 single full-length molecules of pri-miR-17∼92 across all AcIm concentrations. (**Figure 4b,c and Supplementary Figure 10) (Materials and methods)** Short-read RNA structural profiling (SHAPE-MaP) requires on the order of 1,000 reads per nucleotide to estimate positional reactivities by averaging adduct-induced mutation rates over a bulk population of molecules. As a point of reference, median mutation rates from bulk SHAPE-MaP libraries derived from AcIm modified (25mM and 200mM final) pri-miR-17∼92 were 0.03% and 0.1% respectively with mutual 95^th^ percentile rates of 0.96% and 3.41% which corresponds to a median and 95th percentile rate of 0.951 and 32.4 detected adducts per full length read respectively for 200mM AcIm final concentration. Using the single molecule nature of the nanopore data reported here we directly count the number of called modification sites (putative adduct sites) after statistical testing and detection, which for 150mM is approximately 105 median adducts per full length read (**Figure 4d**), corresponding to a 110 fold enhancement over SHAPE-MaP mutation detection rates. We note that this value may be partially inflated due to the distributed nature of modification detection, however even a conservative lower estimate places the median number of called modification sites for nanoSHAPE above the 95th percentile mutation rate for SHAPE-MaP. An additional important consideration concerns how many single molecule reads are required to obtain an optimal structural estimate. To answer this we subsampled full length reads from n = 1 to n = 1000 and calculated the normalized reactivity profile from the per-read current statistical testing as a function of number of reads. (**Figure 4e**) The Spearman’s rank correlation (rho) of the normalized reactivity against the SHAPE-MaP reactivity profile reached 95% of the maximum correlation around 100-200 reads for each AcIm concentration. In the normalized reactivity profile derived from nanoSHAPE we observe less distinctive reactivity features closer to the 5’ end of the pri-miR-17∼92 transcript. To explore this phenomenon we calculated the Spearman’s rank correlation (rho) on a progressively shortened normalized reactivity profile, trimming from the 5’ end. (**Figure 4f**) This procedure revealed a maximum correlation at about 300 nucleotides from the 5’-end of the transcript (rho = 0.53, 150mM AcIm), indicating that the 5’ end of RNAs may not be well resolved by this approach. This may be due to reduced coverage in this region or alternatively less successful completion of RT to the 5’ end due to the presence of AcIm induced adducts which can cause cDNA truncation. Reverse transcription is not strictly required for direct RNA sequencing using the nanopore platform, however the presence of a cDNA is known to stabilize the sequenced RNA strand increasing overall yield and throughput, potentially favoring the recovery of RNA molecules with a full length cDNA annealed.

### Secondary structure prediction of pri-miR-17∼92 using nanoSHAPE and SHAPE-MaP constraints

Finally, we performed secondary structure prediction using nanoSHAPE and SHAPE-MaP normalized reactivities as pseudo-free energy constraints on the RNA folding algorithm. Predicted centroid structures (RNA secondary structure with the minimal base pair distance to all other structures within the Boltzmann ensemble) of the sequence alone, SHAPE-MaP constrained and nanoSHAPE constrained had minimum free energies (Δ*G*_*cent*_) and ensemble diversities (ED) of −298.8 kcal/mol, −417.5 kcal/mol and −299.73 kcal/mol and 152.9, 87.1 and 86.8 respectively. In order to investigate structural similarities between the three centroid structures we performed a pairwise alignment using BEAGLE (BEar Alignment Global and Local). (**Materials and methods**) Comparison of secondary structures using BEAR encoding allows for high level comparison of secondary structural elements such as stems, loops, internal loops, and bulges.(47) Structural similarity and identity to the unconstrained (sequence alone) centroid structural prediction was similar for both nanoSHAPE (72.7%, 62.8%) and SHAPE-MaP (71.5%, 58.5%) however structural similarity and identity between nanoSHAPE and SHAPE-MaP centroid predictions were slightly lower (65%, 50.4%). (**Supplementary Data 2**) Notable differences in secondary structure between the SHAPE-MaP centroid structure and the nanoSHAPE centroid structure include fewer long-range base pairs and larger loop sizes specifically for hairpins 17 and 19a in nanoSHAPE. (**Figure 5a**) Interestingly, in the unconstrained centroid prediction both hairpin 17 and 19a are predicted to have internal bulges (U151 and G435-U437) at the base of their loops towards the 3’ side (45), (**Supplementary Figure 9b**) suggesting that reactivity at these positions combined with the 3’ to 5’ read direction of direct RNA nanopore sequencing may result in proximal detected reactivity among key loop closing base pairs leading secondary structure prediction to predict larger loop sizes. Importantly, centroid structures for both SHAPE-MaP and nanoSHAPE constrained predictions contained all 6 miR hairpins comprising the 17∼92 cluster. In addition to the centroid structure prediction, we performed partition function calculation for RNA structures arising from SHAPE-MaP constrained and nanoSHAPE constrained pri-miR-17∼92 sequences. (**Figure 5b**) Partition function base pair probabilities between all possible nucleotides (i,j) exhibited highly concordant connectivity patterns indicating agreement between the collective predicted structural ensembles. Overall we determine that nanoSHAPE produces reactivity patterns and predicted structural ensembles for the pri-miR-17∼92 sequence consistent with high-throughput sequencing based RNA chemical probing and structural profiling approaches.

**Figure 5.**
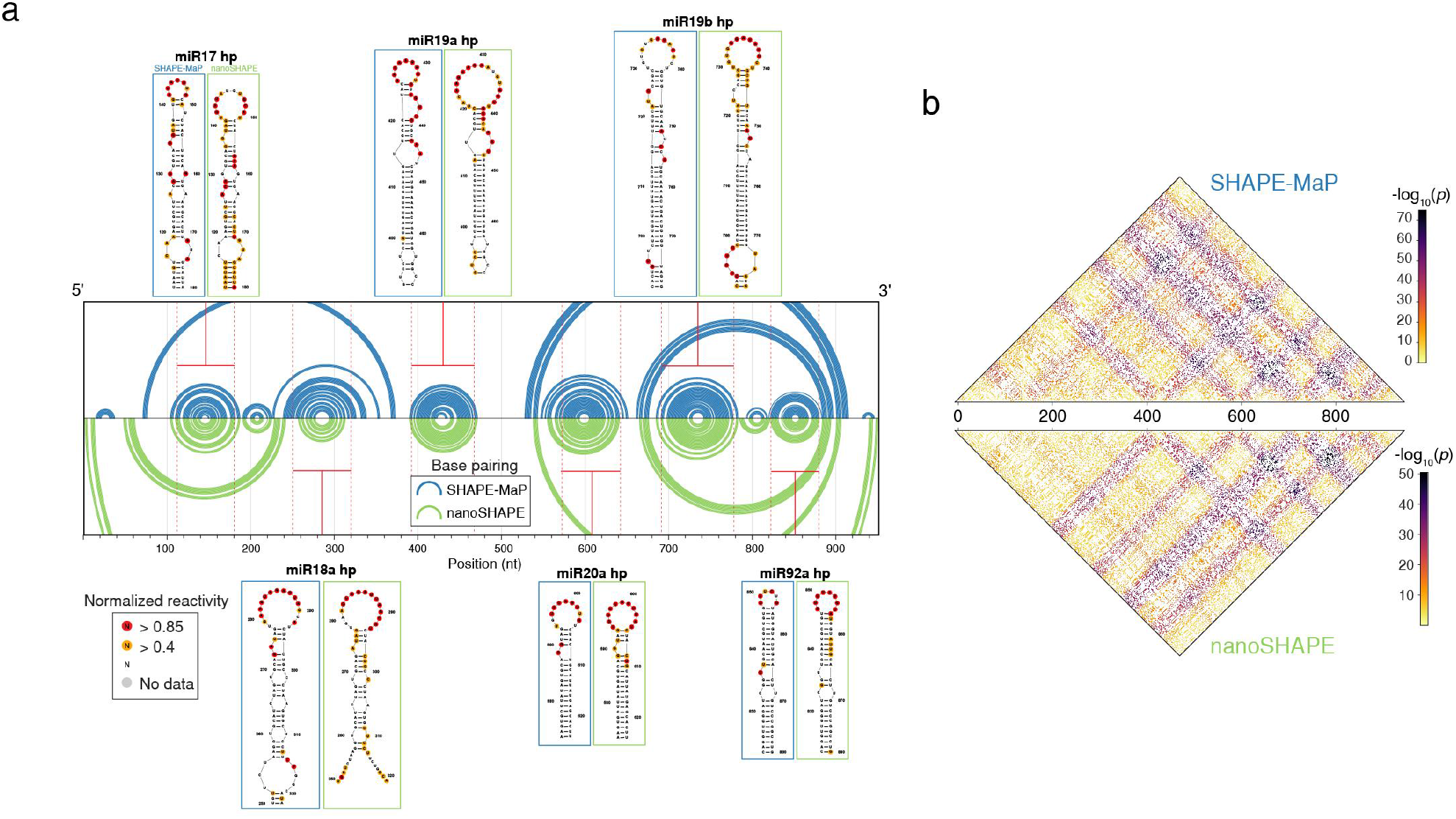
RNA folding comparison using nanoSHAPE reactivities. **a)** Arc diagram comparing the centroid predicted structure for pri-miR-17∼92 RNA using SHAPE-MaP constraints (blue) or nanoSHAPE constraints (green). Secondary structure elements of the constituent miR hairpins are shown with overlaid normalized reactivity from SHAPE-MaP or nanoSHAPE. **e)** Partition function dot plot displaying the probabilities of all possible base pairs i,j in pri-miR-17∼92 using SHAPE-MaP (top) or nanoSHAPE (bottom) reactivity constraints. Probabilities are displayed as - log10(probability base pair (i,j)).

## DISCUSSION

We have demonstrated detection of endogenous RNA modifications and RNA structure probing using the long-read direct RNA nanopore sequencing platform from Oxford Nanopore Technologies. Ribosomal RNA from *E. coli* and *S. cerevisiae* contain numerous and diverse RNA modifications (base, pseudouridine (*Ψ*) and 2’-O-methyl (Nm)) which for the vast majority of positions are stoichiometrically modified, making them ideal systems for benchmarking modification detection using nanopore sequencing. Using in vitro transcribed controls and raw signal to sequence alignment performed by Tombo and Nanopolish, we directly compared current signal distributions identifying rRNA modification positions to within ± 2 nt of known modified sites. We find that both Tombo and Nanopolish discriminate these rRNA modifications comparably.

Further insight into rRNA biogenesis and assembly may be gained by accurate dissection of modification pathways. Additionally, rRNA modifications have been implicated as critical antimicrobial resistance determinants, highlighting the importance of methods for assessing site specific RNA modification patterns.(48) Nanopore sequencing is uniquely positioned to answer questions about the dynamics and ordering of modification installation on rRNA. Nanopore sequencing may ultimately address both accurate quantification and long-range phasing of modifications, but several challenges need to be resolved. Modification quantification is heavily reliant on signal discrimination which is a function of the k-mer sequence context and successful raw signal to sequence alignment. Ultimately development of training sets consisting of known modifications in all possible k-mer sequence contexts will be required. Training sets should allow for the detection and identification of RNA modifications, without the need for IVT control comparison.

In addition to current signal levels we characterized changes in the current level dwell times in direct RNA sequencing. We report that primary sequence and modifications, primarily sugar modifications, can increase dwell times. The sequence dependence of translocation rates has previously been shown for DNA using the Hel308 motor protein and a MspA nanopore (41), but to our knowledge this is the first report corroborating those results on RNA (albeit with an undisclosed motor protein), suggesting broader principles underlying sequence based translocation of nucleic acids. Dwell time changes due to RNA modifications at the pore constriction have been recently reported (49) consistent with our observations. Here we additionally identified that dwell time is dependent on motor protein translocation kinetics mediated at a registration distance (Xr ∼10 nt in the 5’ direction) from the pore constriction. Thus, modifications such as Nm in certain sequence contexts encountering the motor protein manifest their presence as increased dwell times 10 nt in the 3’ direction. Dwell time measurements are partially complicated by the reported dependence of translocation rates on the position of the particular nanopore(s) within the sequencing array. However, we suggest that large dwell times observed in direct RNA nanopore sequencing experiments may be useful to infer sites of Nm or *Ψ* modifications in the absence of IVT controls, provided that the sequence context around the putative modification is not G-rich. Full characterization and incorporation of this extra dimension of information will require nanopore position-specific normalization to faithfully compare translocation rates across the nanopore array.

The ability to detect naturally occurring Nm modifications in rRNA suggested the possibility of detecting other 2’-O adducts which might be introduced exogenously. In an effort to measure RNA structure at the single molecule level we detected adducts on RNA that had been modified with NAI, a common SHAPE reagent. The extremely low read-through rate led us to believe that large bulky adducts are not suitable for translocation through the nanopore, possibly due to unfavorable contacts or steric hindrance between residues of the motor protein and the sugar backbone of the RNA. Searching for a smaller adduct chemical probe we turned our attention to the recently described SHAPE reagent, AcIm.(43) We first demonstrated that AcIm modified RNA is capable of inducing mutational events (Illumina SHAPE-MaP) on par with known SHAPE reagents, establishing AcIm as a genuine SHAPE-MaP reagent. We observed extremely similar reactivity profiles on extracted *E. coli* ribosomes between AcIm, NAI, and NMIA. AcIm has several features of an ideal SHAPE reagent. Its hydrolysis halftime is short, but manageable, which minimizes reaction times and eliminates the need to quench the reagent for in-cell applications. AcIm is highly soluble (up to ∼4.5M in DMSO) which may be advantageous for promoting reaction for challenging experiments, and is commercially available. AcIm is cell and nuclear permeable and, for example, efficiently probes the structure of U1 snRNA. Collectively, for these reasons we expect that AcIm will gain widespread use as a SHAPE/SHAPE-MaP reagent. More work is required to assess whether AcIm may have utility as a single molecule correlated base pair probe (e.g. PAIR-MaP).(50)

We found that the small adduct (2’-O-acetyl) induced by modifying RNA with AcIm is detectable in direct RNA nanopore sequencing experiments, although its presence has the propensity to lower the number of full length reads, overall yield and alignment rates, albeit not as drastically as NAI modified RNA. We used pri-miR-17∼92 as a model system for evaluation of AcIm modification detection due to its well characterized structure comprising multiple hairpins of varying length.

Shifts in positional current signal distributions from nanopore based structural profiling experiments agreed well with the bulk SHAPE-MaP reactivity profile. Using per-read statistical testing against in vitro controls we identified high confidence adduct sites in modified RNAs, finding higher modification rates compared to mutational rates typically encountered in a SHAPE-MaP experiment. Due to higher modification detection rates in nanoSHAPE, optimal correlation with SHAPE-MaP data was obtained after consideration of only ∼200 full length reads, compared to sequencing depth requirements of at least 1k reads per position in SHAPE-MaP for a comparable length target. RNA centroid secondary structural predictions for pri-miR-17∼92 constrained by SHAPE-MaP or nanoSHAPE were very similar. Interestingly, single stranded regions in the centroid structure predicted from nanoSHAPE mostly correspond to low probability base pairing regions in the sequence alone prediction and the SHAPE-MaP constrained prediction suggesting these regions may be dynamic and potentially more susceptible to modification at higher SHAPE reagent concentrations. We do note that nanoSHAPE derived normalized reactivity profiles were less well resolved at the 5’-end compared to SHAPE-MaP. This may be due to a variety of factors including, but not limited to, the intrinsic reactivity characteristics of pri-miR-17∼92, 3’-coverage bias inherent to 3’-to-5’ direct RNA sequencing or incomplete cDNA synthesis on highly modified RNA.

In this preliminary work we intentionally restricted our approach to a well-studied and moderately structured RNA, however it remains to be seen how well nanoSHAPE will perform for RNAs with varying levels of secondary structure. Direct RNA sequencing and by extension nanoSHAPE requires a free 3’-hydroxyl for ligation to the 5’ phosphate of the first sequencing adapter (RTA). We note that electrophilic SHAPE reagents, AcIm included, modify the 3’-hydroxyl, leaving a covalent adduct preventing ligation. We ameliorated this situation by first oxidizing the 3’-end of the RNA followed by a **β**-elimination step leaving a terminal phosphate. Modification at this stage precluded the possibility of 3’ adducts. After modification, the RNA was then treated with a phosphatase to remove the 3’-terminal phosphate before poly(A) tailing and ligation to the RTA adapter. If nanoSHAPE is to be extended transcriptome-wide and to in-cell structural probing experiments, novel methods of library preparation or protection of the 3’-hydroxyl in cells may be required to ensure sufficient yield.

Despite these challenges, nanoSHAPE demonstrates significant promise to dissect complex RNA structures. Long-read single molecule sequencing has strong potential to permit investigation of RNA structural ensembles directly. The high adduct detection rates (i.e. ‘hit-rate’) enabled by direct sequencing of AcIm modified RNA may be crucial to deciphering the RNA energy landscape and alternative folding pathways as well as the phasing of different structural elements, or linking sequence variation to distant structures. Further, we envision a 3-sample experiment (IVT, native, native & AcIm modification) which can be used to investigate native modifications and structure conjointly. Another exciting application of nanoSHAPE is to investigate RNA structure in the context of RNA splicing. Dissecting long-range structural elements that may orchestrate or result from alternative splicing may shed light on regulatory mechanisms underlying this complex phenomena.

Altogether, the technological advancements recently seen in nanopore sequencing have been impressive. We envision improvements in software such as base calling algorithms and signal processing which will benefit the technology markedly. We anticipate further developments to the sequencing chemistry and hardware such as motor proteins, larger nanopore arrays, higher resolution nanopores and improved electronics which will improve yield and throughput and push the limits of sensitivity. As direct RNA nanopore sequencing technology matures and training sets for diverse endogenous and exogenous modifications are developed, this will enable numerous exciting applications which are poised to improve our understanding of fundamental RNA biology.

## DATA AVAILABILITY

Sequence files (multi-Fast5 and summary sequencing files) for all data in this work are available online from NCBI SRA under accession <u>PRJNA634693</u>. (http://www.ncbi.nlm.nih.gov/bioproject/634693)

All computer code is available in the GitHub repository: https://github.com/physnano/rRNA_nanoSHAPE

## SUPPLEMENTARY DATA

Supplementary Data 1 - rRNA modifications

Supplementary Data 2 - BEAGLE structure analysis

## ACKNOWLEDGEMENTS

The authors would like to acknowledge sequencing support from Oxford Nanopore Technologies (NYC). Additionally we would like to thank Marcus Stoiber and Jared T. Simpson for assistance with Tombo and Nanopolish respectively.

## CONFLICT OF INTEREST

W.T. has two patents (8,748,091 and 8,394,584) licensed to ONT. W.S. and W.T. received reimbursement for travel, accommodation and or conference fees to speak at events organized by ONT.

## FUNDING

This work was supported by the National Institutes of Health [HG010538 to W.T. and GM122532 to K.M.W.]

## Notes

http://www.ncbi.nlm.nih.gov/bioproject/634693

