## Supplementary Information for "Direct detection of RNA modifications and structure using single molecule nanopore sequencing"

| Primer name | Primer sequence (5'-3') |
|---|---|
| ec16f_t7p | TAATACGACTCACTATAGGGAAATTGAAGAGTTTGATCATGGCTC |
| ec16r | TAAGGAGGTGATCCAACCGCAGG |
| ec23f_t7p | TAATACGACTCACTATAGGGGGTTAAGCGACTAAGCGTACACGGT |
| ec23r | AAGGTTAAGCCTCACGGTTCATTAG |
| sc18f_t7p | TAATACGACTCACTATAGGGTATCTGGTTGATCCTGCCAGTAGTC |
| sc18r | TAATGATCCTTCCGCAGGTTCACCTAC |
| sc25f_t7p | TAATACGACTCACTATAGGGGTTTGACCTCAAATCAGGTAGGAGTA |
| sc25r | ACAAATCAGACAACAAAGGCTTAATCTC |

**Supplementary Table 1 - Primers for rDNA amplification from *E. coli and S. cerevisiae* gDNA.**

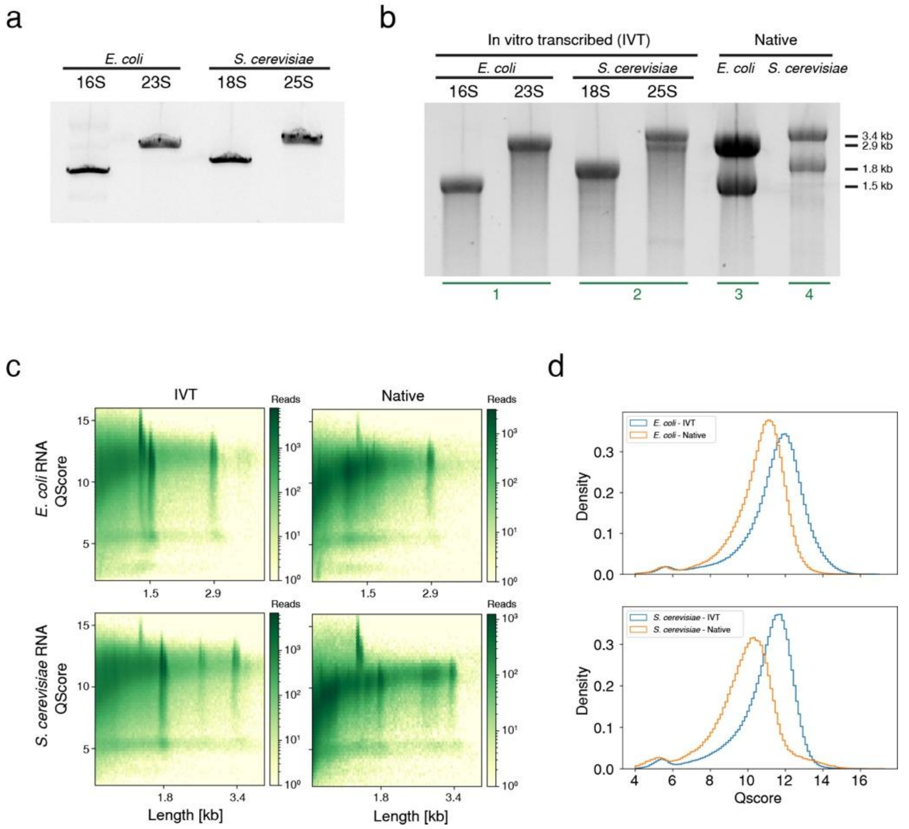


**Supplementary Figure 1 - Generation of IVT controls and QC metrics from nanopore sequencing.** a) SSU and LSU rDNA amplified from *E. coli* and *S. cerevisiae* gDNA with appended 5' T7 promoter. b) In vitro transcription (IVT) of corresponding transcription templates. Extracted total RNA (native) is shown at right. Green lines and numbering indicate the RNA used for individual MinION nanopore experiments. c) Quality score vs read length for IVT and Native samples from *E. coli* and *S. cerevisiae*.

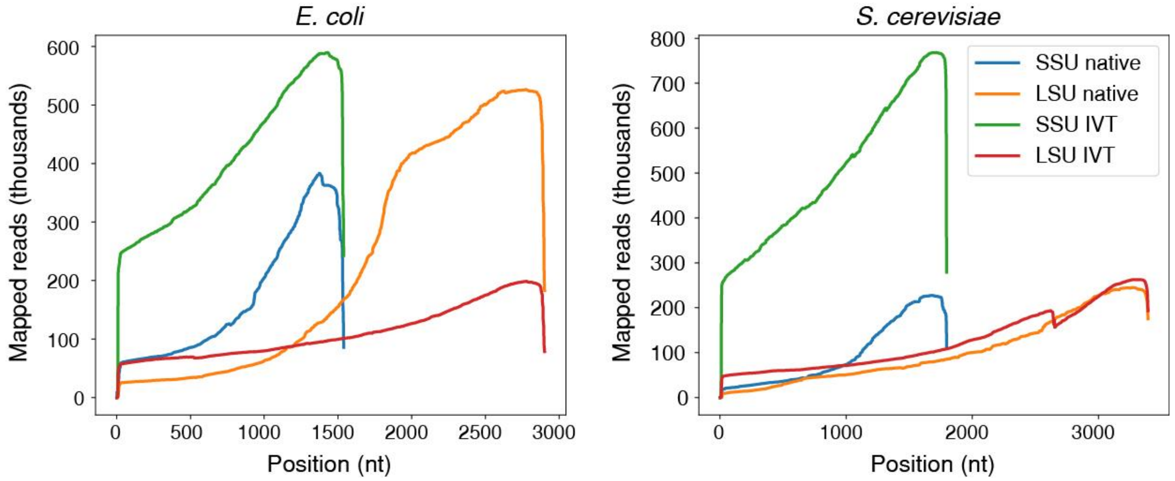


**Supplementary Figure 2 - Mapped rRNA coverage.** Mapped read coverage after processing with *Tombo*.

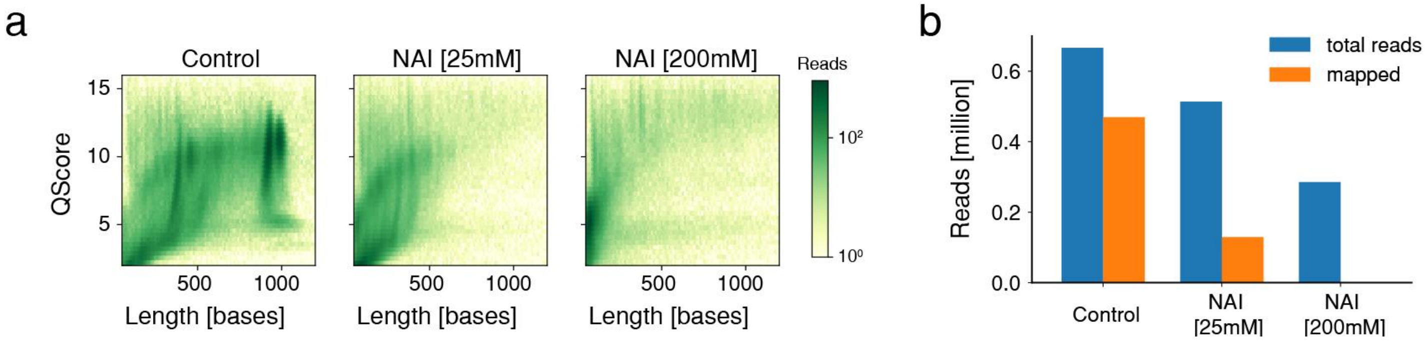


**Supplementary Figure 3 - QC and alignment metrics of NAI modified RNA.** a) Quality score vs read length for control (unmodified) and NAI modified (25mM and 200mM final concentration) of pri-miR 17~92 cluster. b) Number of total reads and mapped reads for control (unmodified) and NAI modified reads.

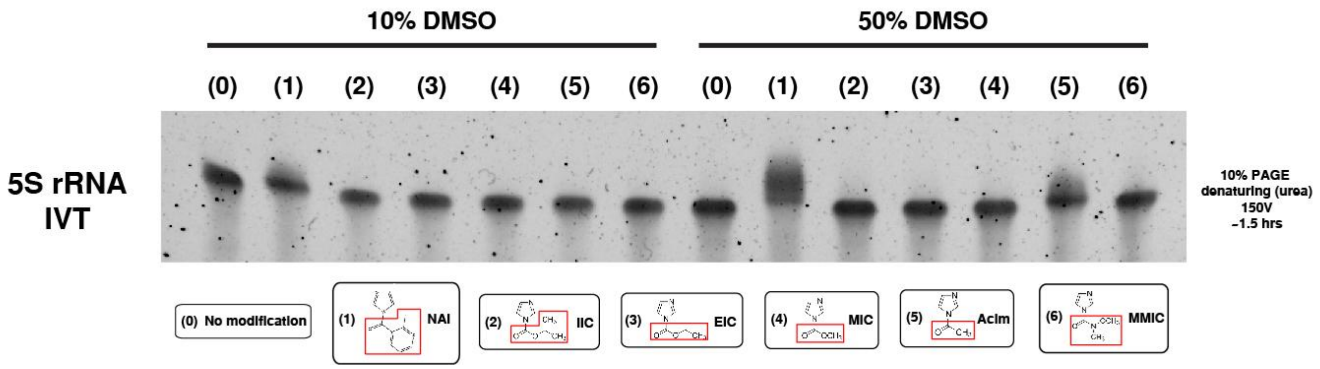


**Supplementary Figure 4 - RNA reactivity for small-adduct RNA-reactive acylation reagents.** 5 putative acylation reagents along with 1 known SHAPE reagent (NAI) were tested for reactivity against RNA in addition to a negative control (DMSO). In vitro transcribed 5S rRNA was modified in non-denaturing (10% DMSO) and denaturing (50% DMSO) conditions for 2 hours prior to resolving the RNA on a denaturing PAGE gel. Chemical modification key:

(0) DMSO vehicle only
(1) NAI (2-methylnicotinic acid imidazolide)
(2) IIC (isopropyl 1H-imidazole-1-carboxylate)
(3) EIC (ethyl imidazole-1-carboxylate)
(4) MIC (methyl imidazole-1-carboxylate)
(5) AcIm (acetylimidazole)
(6) MMIC (N-methoxy-N-methyl-1H-imidazole-1-carboxamide)

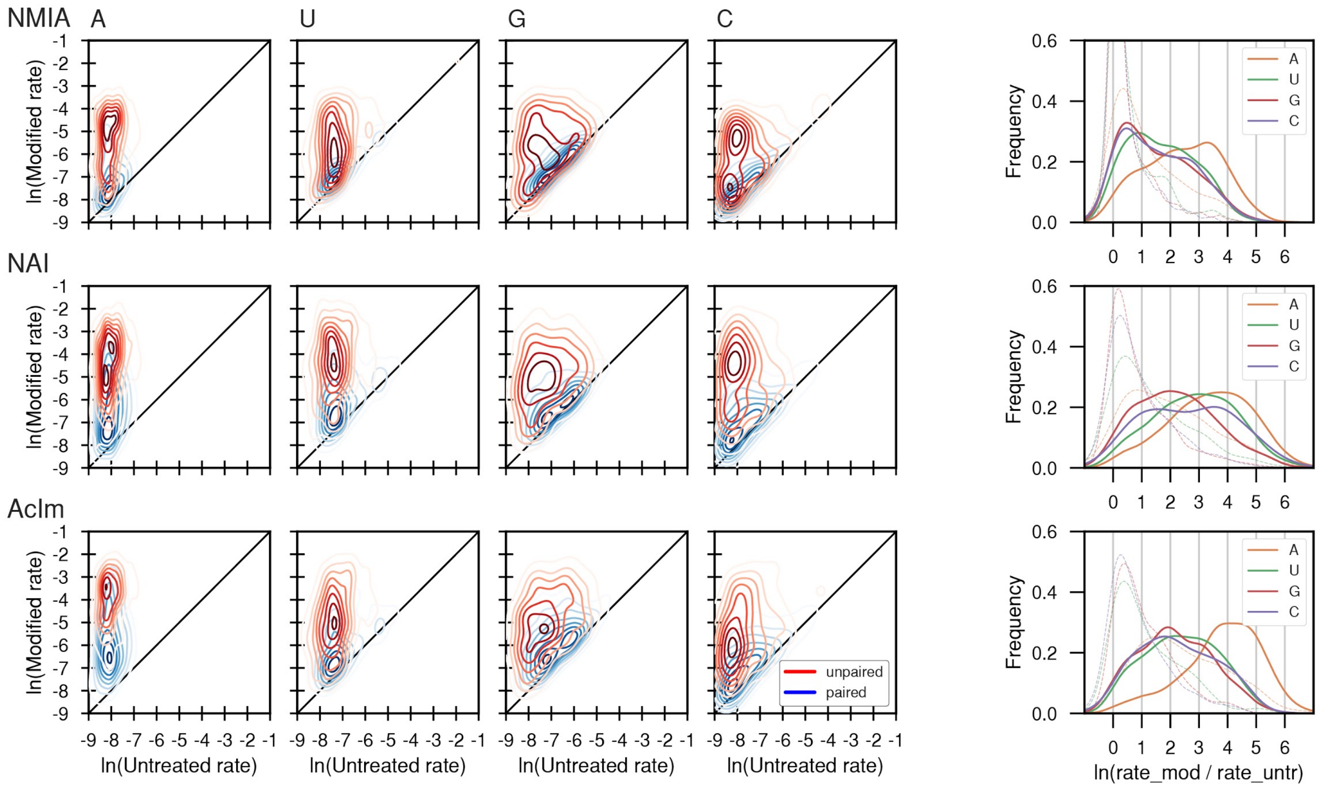


**Supplementary Figure 5. Per-nucleotide MaP mutation rates for SHAPE reagents.** Left four columns show two-dimensional kernel density estimates for mutation rates based on adduct-induced mutation rates for *E. coli* 16S and 23S rRNAs, with unmodified control and SHAPE-modified rates on the x- and y-axes, respectively. Separation between peak density distributions for paired (blue) versus unpaired (red) positions indicates selective reactivity with respect to flexible nucleotides. Right-most column shows one-dimensional kernel density estimates for background-corrected mutation rates, where “rate_mod” is the SHAPE-modified rate and “rate_untr” is the unmodified control rate at a given position. Paired nucleotides are shown with a dashed line, and unpaired nucleotides with a solid line. All three reagents show the highest apparent reactivity with unpaired adenosine residues.

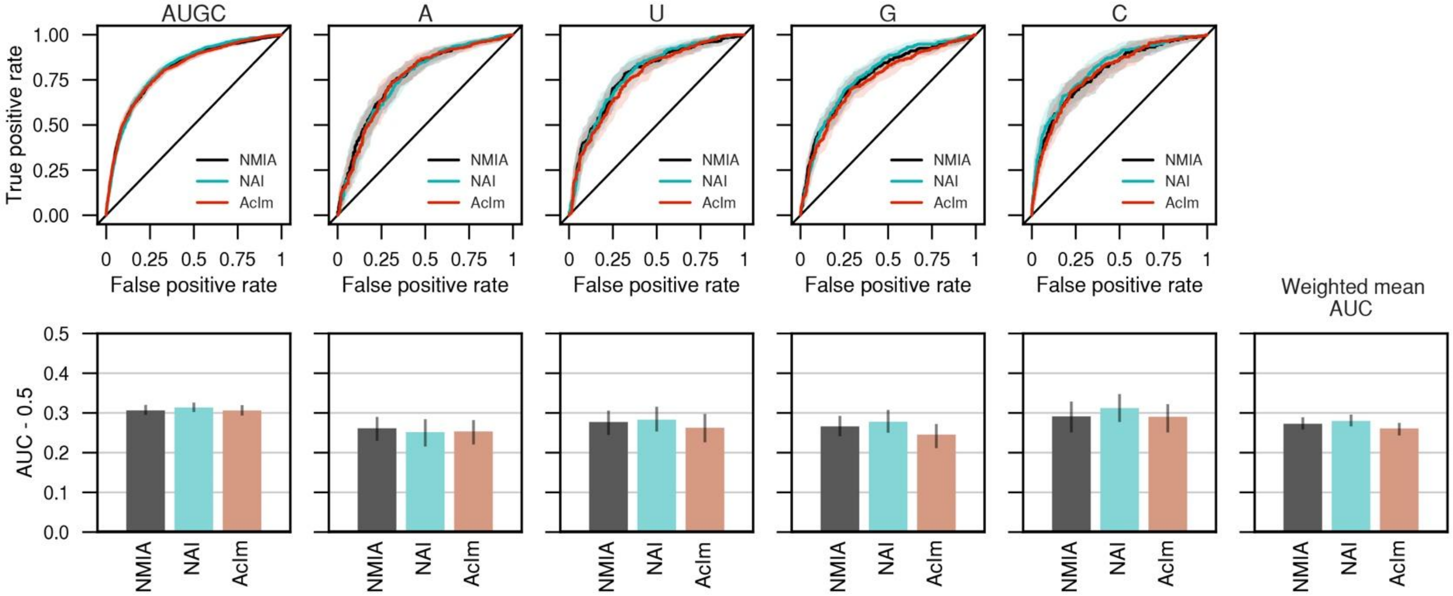


**Supplementary Figure 6 - Receiver operator characteristic (ROC) curves for MaP reactivities as a function of base pairing status for SHAPE reagents.** Analysis is based on reactivities for three reagents using 16S and 23S rRNAs. Left column plots all pooled nucleotides, and right four columns are broken down by each canonical ribonucleotide. True positive rate: unpaired nucleotides with reactivity above a given threshold / total unpaired nucleotides. False positive rate: base-paired nucleotides with reactivity above a given threshold / total base-paired nucleotides. AUC: area under the ROC curve. Shaded regions are 95% bootstrap confidence intervals. Error bars indicate 95% bootstrap confidence intervals on the AUC. All three reagents show comparable agreement with base pairing status across all four ribonucleotides.

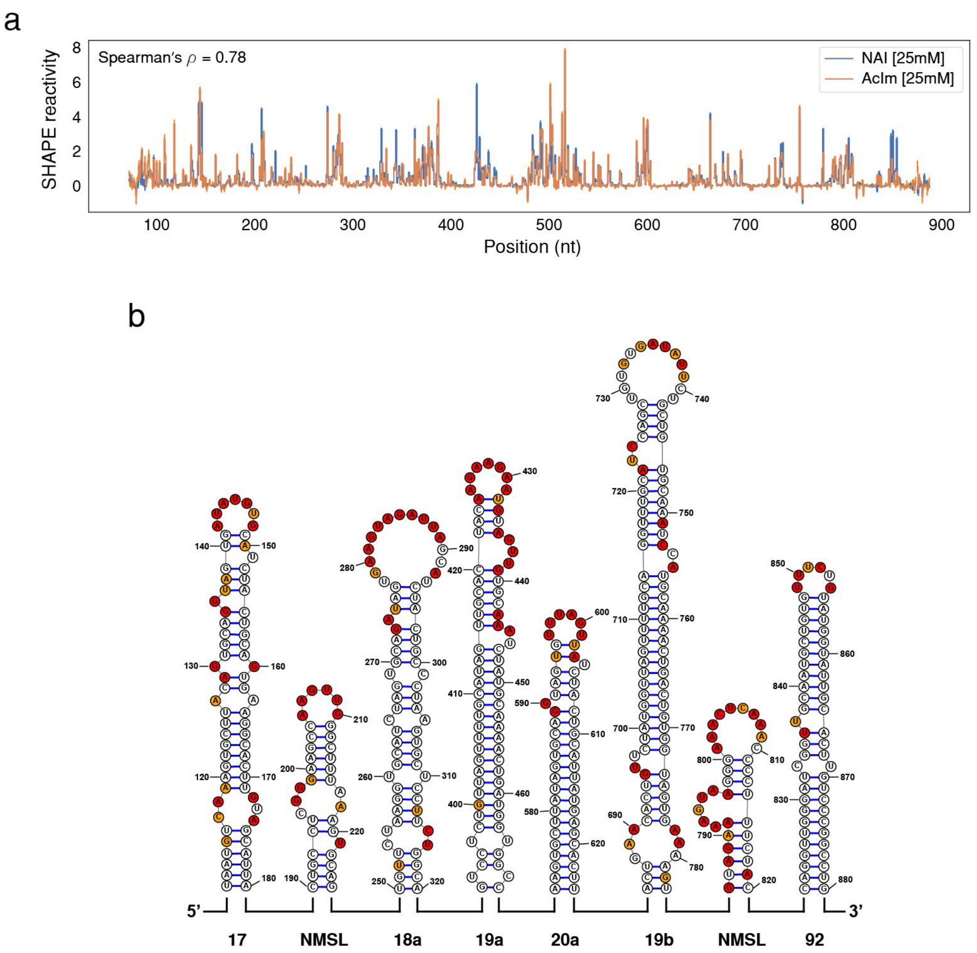


**Supplementary Figure 7 - SHAPE-MaP profile of pri-miR 17~92. a)** Normalized reactivity profiles for pri-miR 17~92 using NAI and AcIm both at 25mM final concentration. Error bars are normalized standard deviation. **b)** Secondary structure of the pri-miR 17~92 transcript (miR hairpins shown) with the AcIm SHAPE-MaP reactivity overlaid. [REF to structure?]

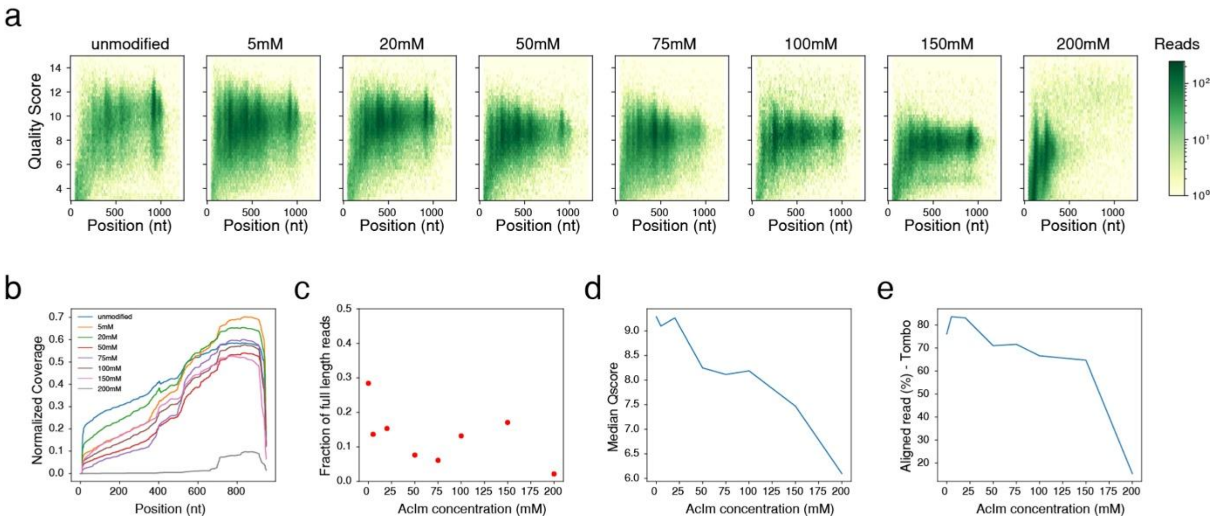


**Supplementary Figure 8 - QC metrics direct RNA sequencing (flongle flow cell) of AcIm modified pri-miR-17~92.** a) Quality score and read length heatmaps for pri-miR-17~92 modified with 0 (unmodified control), 5, 20, 50, 75, 100, 150, and 200mM final concentration of AcIm. b) Read coverage for AcIm modified pri-miR-17~92. c) Fraction of full length reads obtained for AcIm modified pri-miR-17~92. d) Median Qscore for AcIm modified pri-miR 17~92. e) Aligned read percentage after raw signal alignment with sequence using Tombo for AcIm modified pri-miR-17~92.

| Concentration AcIm [mM] | Reads | Full length reads (fastq) | Median Qscore | Tombo alignment [%] | Total number aligned full length reads (Tombo) |
|---|---|---|---|---|---|
| 0 | 68693 | 17216 | 9.29 | 76.1 | 13101 |
| 5 | 163000 | 21173 | 9.1 | 83.6 | 17701 |
| 20 | 117546 | 16939 | 9.26 | 83.1 | 14076 |
| 50 | 97344 | 6720 | 8.25 | 71 | 4771 |
| 75 | 120932 | 6944 | 8.11 | 71.6 | 4972 |
| 100 | 52947 | 6389 | 8.19 | 66.6 | 4255 |
| 150 | 77992 | 12182 | 7.5 | 64.7 | 7882 |
| 200 | 108096 | 1669 | 6.1 | 15.4 | 257 |

**Supplementary Table 1 - Flongle runs read summary**

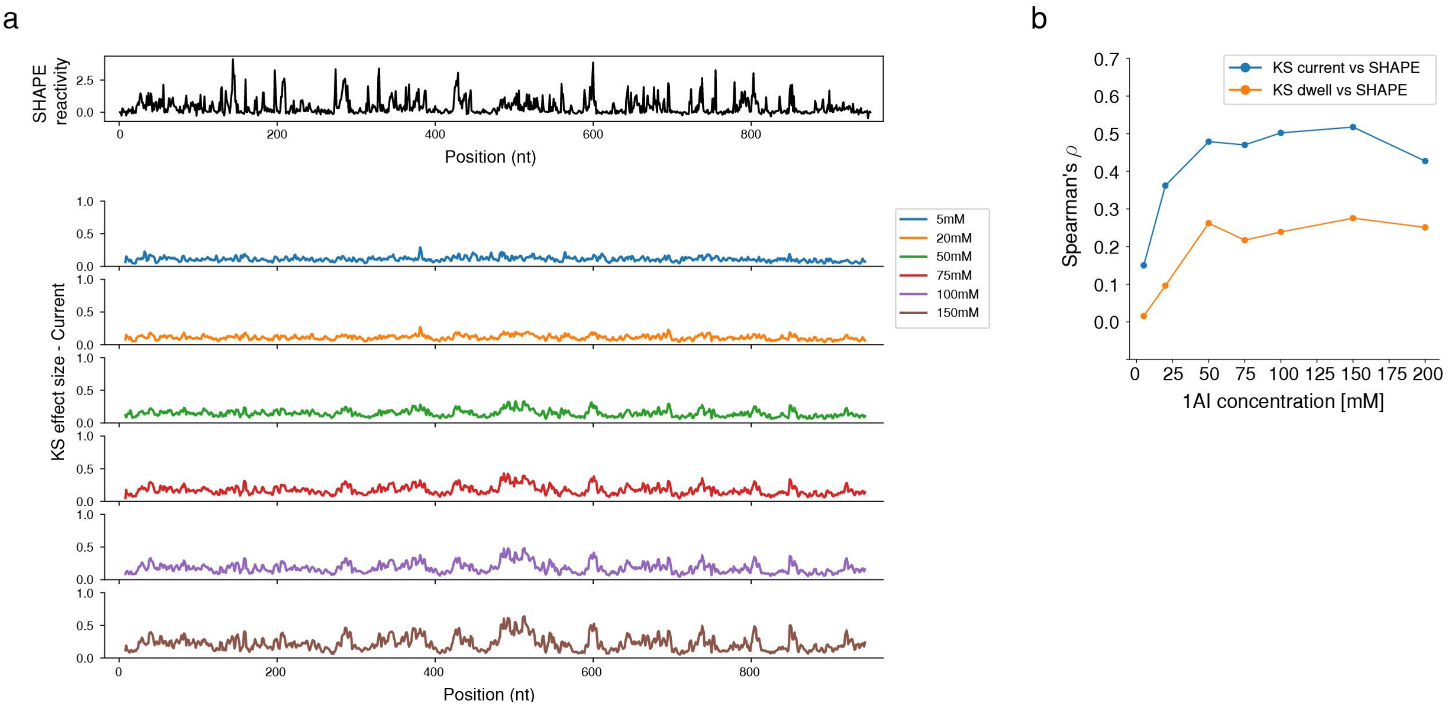


**Supplementary Figure 9 - KS profiles of AcIm modified RNA. a)** KS profiles for AcIm modified pri-miR-17~92 RNA. **b)** Spearman's rho correlation of KS profile for current and dwell time against the SHAPE-MaP for the AcIm modification concentrations.

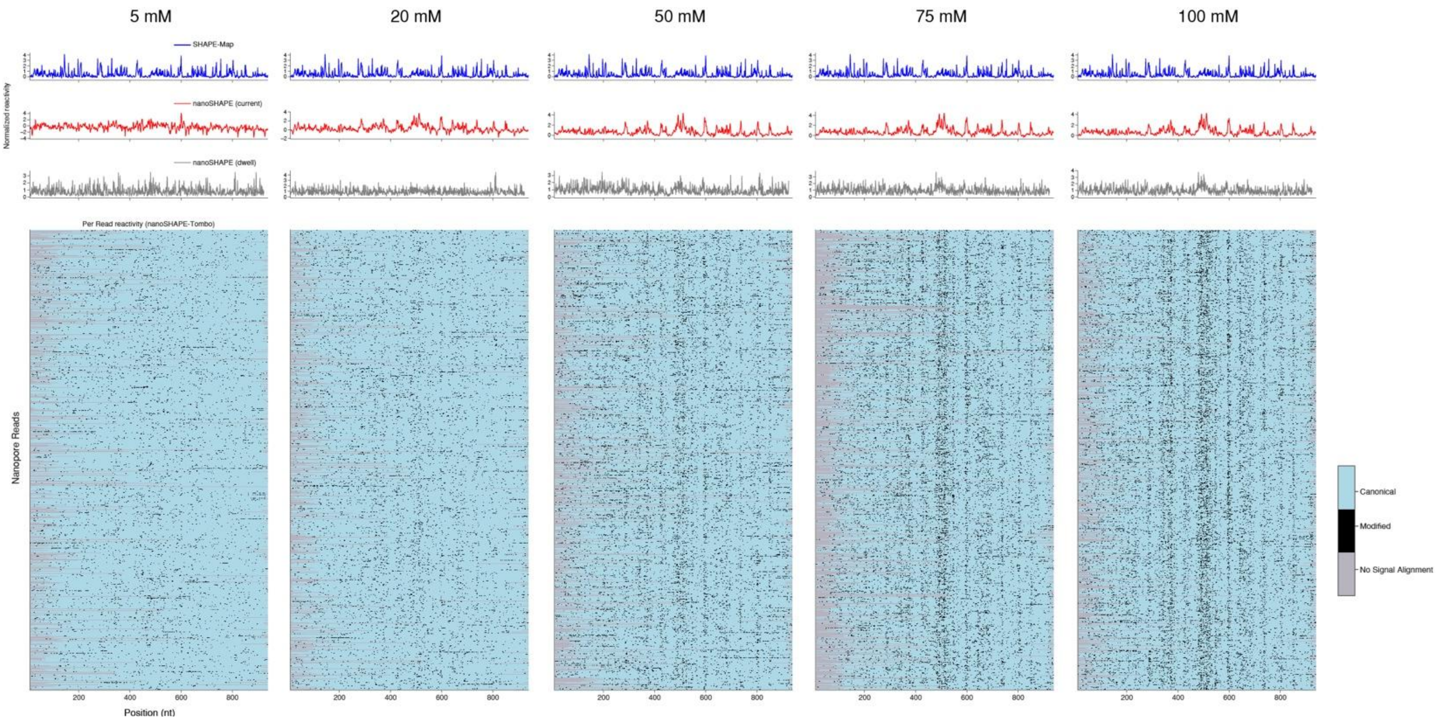


**Supplementary Figure 10 - Per read statistical testing and normalized reactivity profiles across AcIm concentrations (5 mM, 20 mM, 50 mM, 75 mM, 100 mM)**
